# Acoustic phenotyping reveals sublethal pesticide effects in bees using BEEhaviourLab

**DOI:** 10.64898/2026.03.03.709383

**Authors:** Rachel H. Parkinson, Oliver N. F. King, Jung Chun (Zaza) Kuo, Kieran Walter, Ash Silva, Jennifer Scott, Cait Newport, Geraldine A. Wright, Stephen Roberts

## Abstract

Behavioural changes can provide early indicators of physiological stress, yet remain difficult to quantify at scale. Acoustic behaviours such as buzzing are widespread in insects but are rarely incorporated into automated ecotoxicological assays. We developed BEEhaviourLab, a multimodal platform combining tag-free video tracking with automated acoustic phenotyping, and validated it across four insect pollinator species (honey bee, *Apis mellifera*; bumble bee, *Bombus terrestris*; ivy bee, *Colletes hederae*; and hoverfly, *Eristalis tenax*). A shared computer-vision model accurately detected diverse taxa, and automated acoustic estimates closely tracked manually annotated variation in buzzing activity. Locomotor and acoustic activity were strongly associated but not redundant: buzzing also occurred during periods of low locomotor activity, capturing behavioural variation that video alone would miss. We then used BEEhaviourLab to quantify effects of the veterinary antiparasitic moxidectin in the bumble bee, *B. terrestris*. Acute contact exposure caused dose-dependent reductions in movement probability, locomotor speed and buzzing, alongside altered diel activity, highlighting potential risks from veterinary pharmaceuticals to pollinators. Integrating movement probability, locomotor speed and buzzing probability identified an estimated behavioural-effect threshold of 2.73 ng bee⁻¹, below the lowest tested dose associated with significantly elevated mortality hazard (78.2 ng bee⁻¹). These findings demonstrate that acoustic phenotyping adds complementary information to automated video tracking and that multimodal behavioural monitoring can reveal toxicant effects well before conventional survival endpoints, providing a scalable approach for insect ecotoxicology.

## Introduction

Behaviour often provides the earliest detectable signs of physiological disruption^1,2^, yet quantifying behavioural change at scale remains challenging^3^. In toxicology and environmental risk assessment, most regulatory tests still rely on lethal dose estimates obtained from a small number of model species^4^. These assays provide reproducible benchmarks for chemical hazard but reveal little about how chemicals alter behaviours that determine fitness and reproduction, leaving many ecologically relevant effects undetected^5^. This gap largely reflects the difficulty of measuring behaviour in a standardised and scalable way: behavioural responses are inherently variable and multidimensional, making them difficult to translate into quantitative endpoints that can be compared across individuals, experiments, and species^6^.

Among behavioural endpoints, locomotion provides a particularly informative indicator of neurophysiological stress^7,8^. Many neurotoxic compounds disrupt sensory processing or motor control, leading to measurable changes in activity, coordination, or movement patterns^9,10^. Sublethal exposure can translate into large fitness consequences by limiting foraging efficiency, navigation, and predator avoidance^11–13^. Locomotion in flying insects is tightly linked to thoracic flight muscle activity, which also generates acoustic signals through vibration of the thorax and wings. In bees, these vibrations produce characteristic buzzing sounds that accompany locomotion, flight preparation, thermoregulation, defensive signalling, and floral sonication^14–16^. Acoustic behaviour has been particularly well studied in honey bees, where vibrational and airborne signals additionally play roles in communication^17^ and have been used to infer colony-level state, including the prediction of swarming events^18,19^.

Acoustic signals reflect motor output and neuromuscular function and therefore have the potential to provide behavioural indicators of physiological stress while enabling computationally simple and scalable measurement. To test this, we developed BEEhaviourLab, a low-cost platform that combines synchronised video and acoustic recording with automated behavioural analysis. The system supports long-duration experiments across many individuals without physical tags or manual scoring, enabling quantitative comparison of locomotor and acoustic activity under controlled exposure conditions.

As a model case, we examined the effects of moxidectin, a widely used veterinary antiparasitic belonging to the macrocyclic lactone class. Moxidectin is excreted in livestock faeces and can persist in dung for several months, with rainfall facilitating its movement into the surrounding environment^20,21^. Additional dispersal may occur through airborne particulate matter, with moxidectin detected in dust collected near cattle feedlots^22^. Experimental studies have also demonstrated that moxidectin can persist in soil and be taken up into above-ground plant tissues^23^. These pathways create opportunities for non-target insect exposure through contact with contaminated soil, vegetation or airborne particles, including among ground-nesting bees and species that collect mud or other materials for nest construction. Indeed, moxidectin has been detected in nesting materials and developing insects collected near cattle feedlots, providing direct evidence of environmental exposure among cavity-nesting insect assemblages^24^. However, exposure levels in ground-nesting bumble bees remain unknown.

Macrocyclic lactones act on glutamate– and γ-aminobutyric acid (GABA)-gated chloride channels in invertebrates^25^, suppressing synaptic transmission and disrupting behaviours linked to sensory processing and motor control^20,26–28^. Laboratory studies have demonstrated lethal and sublethal effects of macrocyclic lactones in honey bees and mason bees^29,30^. However, despite evidence of environmental exposure and toxicity, their effects on behaviours that underpin pollination, including movement and daily activity patterns, remain poorly characterised in bees.

Using BEEhaviourLab, we quantified insect activity using synchronised video and acoustic measurements across four pollinator species selected to represent variation in body size, morphology, social organisation and flight dynamics. Video-based tracking captured clear differences in locomotor and diel behaviour in honey bees (*Apis mellifera*), bumble bees (*Bombus terrestris*), ivy bees (*Colletes hederae*), and hoverflies (*Eristalis tenax*). Acoustic behaviour proved particularly informative in bees, especially the eusocial species *A. mellifera* and *B. terrestris*, which produced robust buzzing activity. In bumble bees, acoustic measures of buzzing also detected behavioural disruption following moxidectin exposure at doses comparable to those detected by video-based locomotion metrics. Together, these results demonstrate that BEEhaviourLab enables scalable multimodal behavioural phenotyping across insect taxa and identify acoustic behaviour as a powerful and previously underused indicator of diel behaviour and stress in eusocial bees.

## Results

### A single multi-species detector enables accurate, tag-free tracking

We evaluated whether a single computer-vision detector could support tag-free tracking across diverse pollinator taxa (Figure 1). We selected four ecologically relevant and experimentally tractable insect pollinators spanning gradients in body size, morphology, social organisation (eusocial and solitary), and flight dynamics: *Bombus terrestris* (eusocial bumble bee; female worker body length range 11–17 mm^31^), *Apis mellifera* (eusocial honey bee; 12–15 mm^32^), *Colletes hederae* (solitary bee; 11.5–14.5 mm^33^), and *Eristalis tenax* (solitary hoverfly with a distinct dipteran body plan and flight kinematics; 14–16 mm^34^).

**Figure 1:**
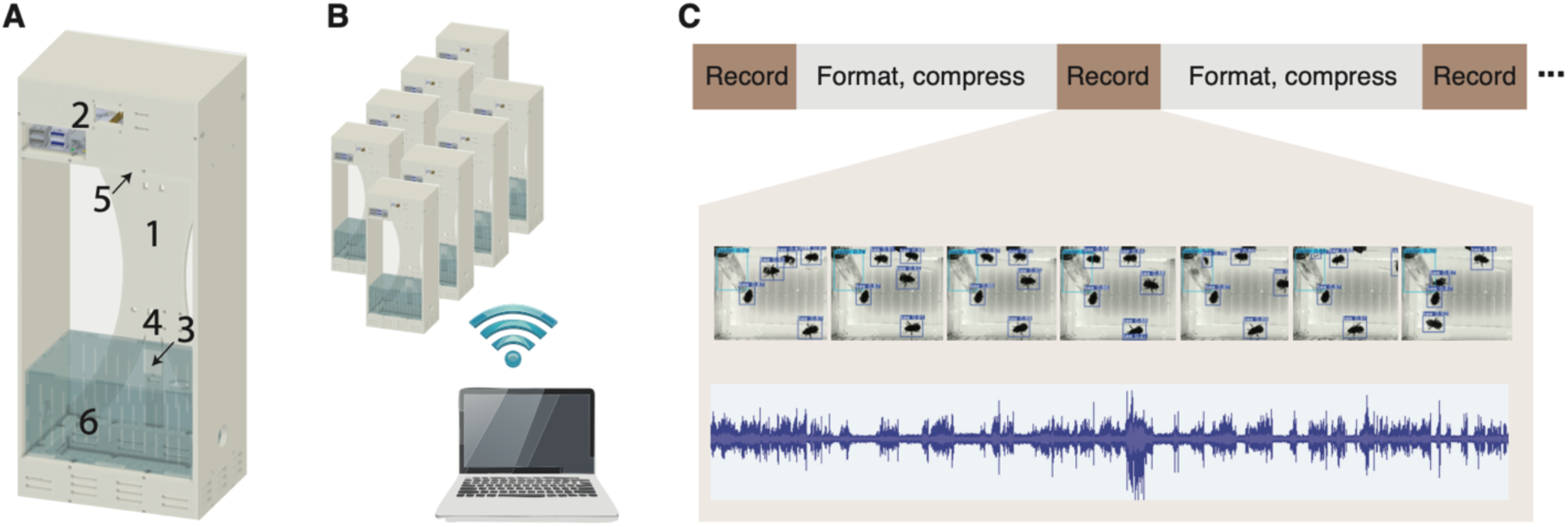
High-throughput insect behavioural tracking using BEEhaviourLab. A: Schematic of a single BEEhaviourLab unit, showing the Perspex enclosure (1), housing a Raspberry Pi (2), temperature and humidity probe (3), microphone (4), and camera (5). Insects are housed within a transparent recording cage (6). B: Multiple units operate in parallel and communicate wirelessly with a central control computer. C: User-defined recording protocols enable flexible specification of recording duration (brown), inter-recording intervals (grey), and number of recording segments. During inter-recording intervals, video, audio, sensor data, and experimental metadata are formatted, compressed, and transferred to the central computer via Wi-Fi.

A YOLOv8n model trained exclusively on *B. terrestris* images (model 1, 155 annotated images) achieved high detection performance for bumble bees (precision, recall, and F1 > 0.97; Table 1) but generalised poorly to other taxa. Performance was lowest for *C. hederae* (precision and recall < 0.50) and was moderately reduced for *A. mellifera* and *E. tenax*, particularly under motion blur. We therefore trained a second model (model 2) using 455 annotated images from all four species (bumble bee: 155 images; other species: 100 images). This multi-species model achieved consistently high precision, recall, F1-score and mAP50 across taxa (mAP50 ≥ 0.98; Table 1), although localisation performance at more stringent overlap thresholds was lower (mAP50–95 = 0.64–0.85). Performance remained high when the final model was evaluated on the independent test dataset comprising recordings excluded from both training and validation (mAP50 = 0.98; Table 1). Together, these results show that incorporating morphological diversity during training enabled a single lightweight detector to generalise across pollinator taxa within the shared experimental environment.

**Table 1:** YOLO model benchmarking across pollinator species.

| **Model** | **Species** | **N validation images** | **Precision** | **Recall** | **F1-Score** | **mAP50** | **mAP50-95** |
|---|---|---|---|---|---|---|---|
| model 1 | bumble bee | 50 | 0.98 | 0.97 | 0.98 | 0.98 | 0.77 |
|  | ivy bee | 38 | 0.50 | 0.45 | 0.47 | 0.47 | 0.37 |
|  | honey bee | 40 | 0.76 | 0.67 | 0.74 | 0.74 | 0.39 |
|  | hoverfly | 38 | 0.83 | 0.67 | 0.75 | 0.75 | 0.36 |
| model 2 | bumble bee | 50 | 0.98 | 0.98 | 0.98 | 0.98 | 0.85 |
|  | ivy bee | 38 | 0.99 | 0.99 | 0.99 | 0.99 | 0.85 |
|  | honey bee | 40 | 1.00 | 1.00 | 1.00 | 1.00 | 0.84 |
|  | hoverfly | 38 | 1.00 | 0.99 | 0.99 | 0.99 | 0.64 |
|  |  | **N test images** |  |  |  |  |  |
| model 2 | bumble bee | 20 | 0.98 | 0.97 | 0.97 | 0.98 | 0.84 |
|  | honey bee | 20 | 0.99 | 0.98 | 0.98 | 0.98 | 0.82 |

### Automated acoustic detection reliably captures variation in buzzing activity

Manual annotation of 50 recordings identified 372 buzzing or flight-sound intervals, totalling 353.9 s (5.90 min) of acoustic activity. Individual bouts were generally short (median = 0.26 s; mean = 0.95 s), and 23 recordings contained no manually detected buzzing. At the clip level, automated estimates of the proportion of time buzzing were strongly correlated with manual annotations (Pearson’s r = 0.91; Spearman’s ρ = 0.89), with similarly strong agreement for B. terrestris (r = 0.93) and the other species combined (r = 0.87). Mean absolute error was 0.088.

The detector was highly sensitive at the frame level, detecting 98% of manually labelled buzzing frames (recall = 0.98), with specificity of 0.89 and balanced accuracy of 0.94. Precision was lower (0.50; F1 = 0.66), reflecting a tendency to classify additional frames as buzzing rather than to miss labelled events. Consistent with this, the automated estimate of the proportion of time buzzing was higher on average than the manual estimate (0.20 versus 0.12; mean difference = +0.079). False-positive detections occurred predominantly in recordings containing genuine buzzing (96.3% of all false-positive frames), rather than in completely silent recordings (3.7%). Furthermore, the false-positive rate among manually silent frames was substantially lower in completely silent recordings (0.8%) than in recordings containing genuine buzzing (21.7%). Thus, although the detector systematically overestimated the absolute duration of buzzing, false detections were primarily associated with recordings containing genuine acoustic activity rather than occurring in otherwise silent recordings.

### Multimodal tracking reveals species-specific activity patterns

Using the multi-species detector, we quantified locomotor activity over 48 h (Figure 2A). Speeds were strongly bimodal, so insects were classified as active above a threshold of 0.95 px s⁻¹ (Figure S1, see Methods). Locomotion was analysed using a two-part mixed-effects framework that separates the probability of movement from movement speed conditional on activity. Species differed markedly in both the probability of movement and locomotion speed when active. Bumble bees showed the highest probability of movement (P = 0.76, 95% CI 0.68–0.82), whereas honey bees had the highest average speeds when active (2.16 mm s⁻¹, 95% CI 1.92–2.43). Hoverflies were both less likely to move and slower when active. Locomotor activity declined slightly over time, but this effect was small relative to interspecific differences. Activity also showed strong 24-h periodicity, with species significantly modifying both harmonic components (sin24: χ² = 58.0, p < 0.0001; cos24: χ² = 162.1, p < 0.0001). Honey bees and ivy bees showed high-amplitude diurnal rhythms, whereas bumble bees remained more active across the diel cycle, producing weaker diel patterns. Hoverflies showed comparatively weak and irregular diel structure.

**Figure 2:**
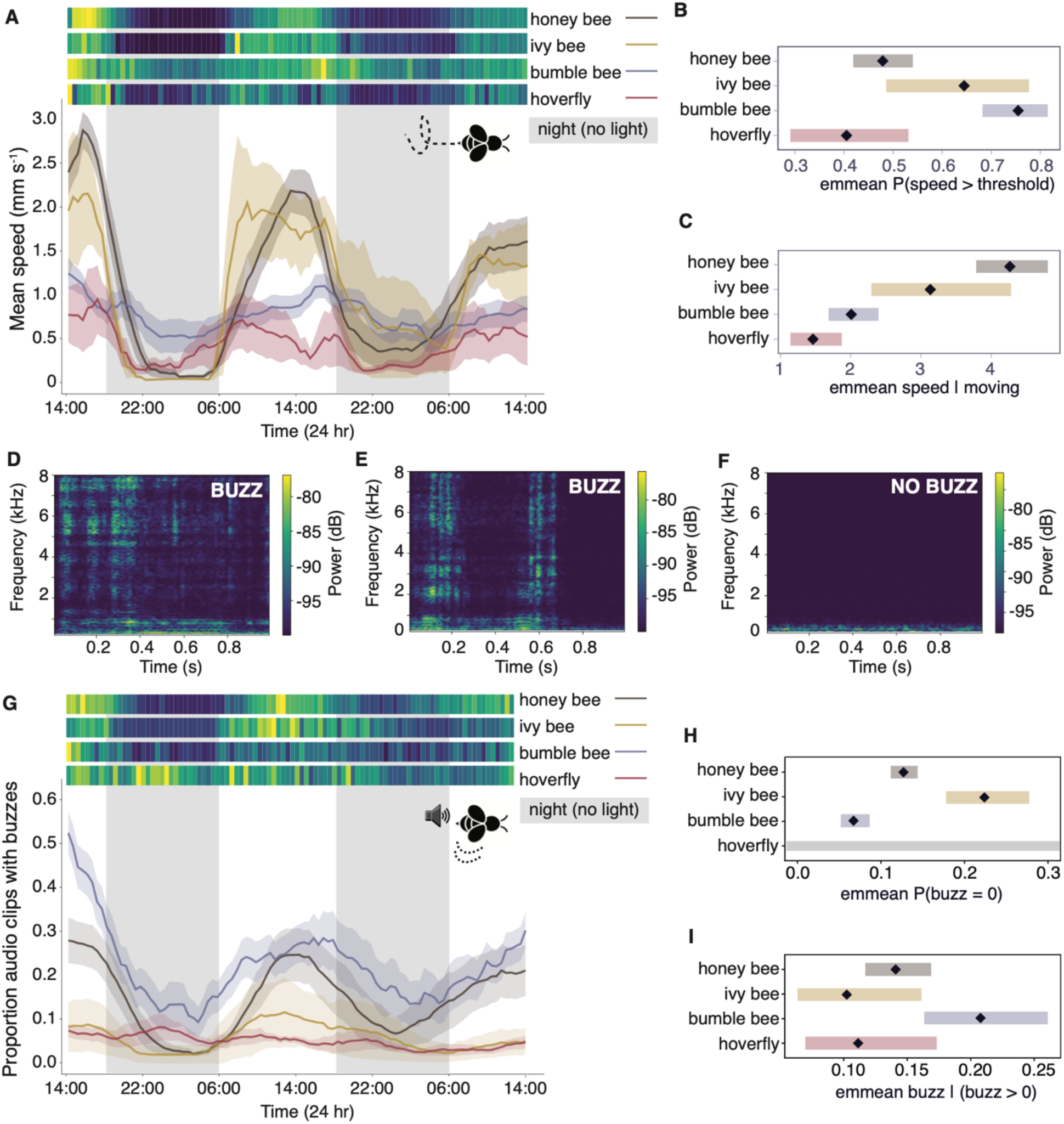
Cross-species locomotor and acoustic activity patterns. A: Locomotion speed over 48 h by species; heatmaps are scaled to species-specific maxima, with lines showing mean ± SD across individuals. B: Estimated marginal means (EMMs) for the probability of movement (speed above the movement threshold). C: EMMs for locomotion speed conditional on movement. D–F: Example spectrograms showing audio clips with buzzing (D–E) or no buzzing (F). G: Acoustic activity over 48 h by species, expressed as the proportion of each 60 s recording containing detected buzzing-related sound; heatmaps are scaled to species-specific maxima and lines show mean ± SD. H: EMMs for the proportion of time buzzing conditional on non-zero acoustic activity. I: EMMs for proportion of clips with buzzes.

Acoustic activity also differed between species (Figure 2G). For each recording, acoustic activity was quantified as the proportion of the 60 s recording classified as containing buzzing-related sound. Bumble bees showed the highest levels of acoustic activity and ivy bees the lowest, whereas hoverflies produced short, relatively frequent but unsustained buzzes. Among recordings containing buzzing (Figure 2H), bumble bees exhibited the highest mean activity (0.208, 95% CI 0.163–0.261), significantly exceeding ivy bees (0.102, 95% CI 0.063–0.161; Holm-adjusted p = 0.039, EMM) but not hoverflies (0.111, 95% CI 0.070–0.173; p = 0.058, EMM). Honey bees showed intermediate activity (0.141, 95% CI 0.117–0.169) and did not differ significantly from the other species after correction.

Species also differed in the model-estimated probability that an entire recording contained no buzzing (Figure 2I). Bumble bees had a lower probability of complete inactivity (0.067, 95% CI 0.052–0.086) than honey bees (0.127, 95% CI 0.112–0.144; p < 0.001) and ivy bees (0.224, 95% CI 0.178–0.278; p < 0.001). Zero-inflation estimates for hoverflies were unstable because no hoverfly recording lacked detected buzzing, reflecting their frequent but brief buzzes. Across species, acoustic activity increased modestly over time (β = 0.071 ± 0.018 SE, p < 0.001).

### Moxidectin exposure induces dose-dependent behavioural impairment in bumble bees

We next examined the effects of acute moxidectin exposure on bumble bee behaviour over 48 h following contact exposure to moxidectin or the vehicle control (Figure 3). Increasing dose produced a strong, monotonic reduction in the probability of movement (log-dose effect: β = −0.91 ± 0.12 SE, z = −7.58, p < 0.0001), declining from 0.66 in controls to 0.50 at the lowest dose (0.782 ng bee^-1^) and to 0.13 at the highest dose (782 ng bee^-1^, Figure 3B). Among bees that were active, locomotion speed also decreased with increasing dose (β = −0.13 ± 0.03 SE, z = −4.49, p < 0.0001), but the magnitude of this effect was smaller (Figure 3C).

**Figure 3:**
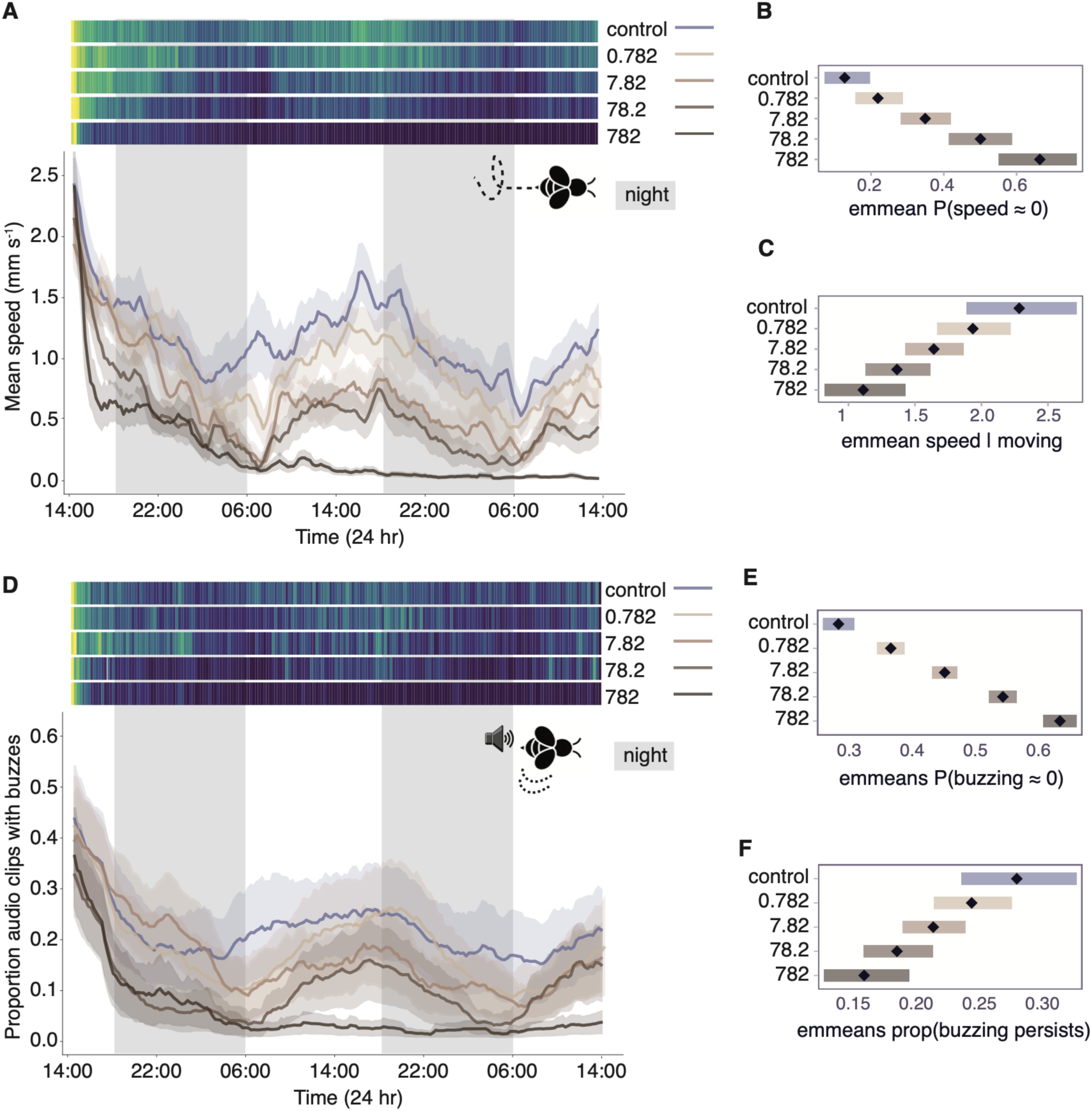
Moxidectin induces dose-dependent reductions in locomotion and buzzing. A: Mean locomotion speed over 48 h across moxidectin treatments; heatmaps are scaled to the maximum within each treatment group. B: Estimated marginal means (EMMs) for the probability that speed falls below the movement threshold, showing dose-dependent increases in inactivity. C: EMMs for locomotion speed conditional on movement, indicating more modest dose-dependent reductions among active bees. D: Mean proportion of time containing detected buzzing-related sound within each 60 s recording over 48 h across treatments; heatmaps are scaled to treatment-specific maxima. E: EMMs for the probability that an entire recording contained no detected buzzing, showing dose-dependent increases in complete acoustic inactivity. F: EMMs for proportion of time buzzing conditional on non-zero acoustic activity, showing more modest dose-dependent reductions in recordings where buzzing occurred.

Acoustic measurements showed parallel dose-dependent effects on buzzing behaviour (Figure 3D). Increasing dose increased the probability that an entire recording contained no buzzing activity (zero-inflation component; Figure 3E; β = 0.514 ± 0.022 SE, z = 23.7, p < 0.0001), with model-predicted acoustic inactivity increasing progressively across the dose range. Among recordings containing buzzing, activity declined more modestly with increasing dose (Figure 3F; β = −0.232 ± 0.057 SE, z = −4.10, p < 0.0001) and over time (β = −0.0495 ± 0.0144 SE, z = −3.43, p < 0.001).

Moxidectin exposure also altered the temporal organisation of acoustic activity. Dose significantly modified both 24-h harmonic terms (sin24: χ² = 56.3, p < 0.0001; cos24: χ² = 36.1, p < 0.0001), indicating dose-dependent changes in the shape and timing of diel activity patterns. The lowest dose produced little detectable change, whereas progressively larger alterations were evident at higher doses.

### Buzzing frequently accompanies locomotion but also occurs at low locomotor activity

Across 10,130 recordings from the moxidectin experiment, buzzing and locomotion were strongly associated but not completely coupled (Figure 4A). Buzzing was present in 88.3% of recordings with cage-mean speed above the locomotion threshold, compared with 28.9% of recordings below the threshold. The most common categories were buzzing while moving (42.3%) and silence at low speed (37.1%). Buzzing without substantial locomotion nevertheless accounted for 15.1% of all recordings, while only 5.6% showed locomotion without buzzing.

**Figure 4:**
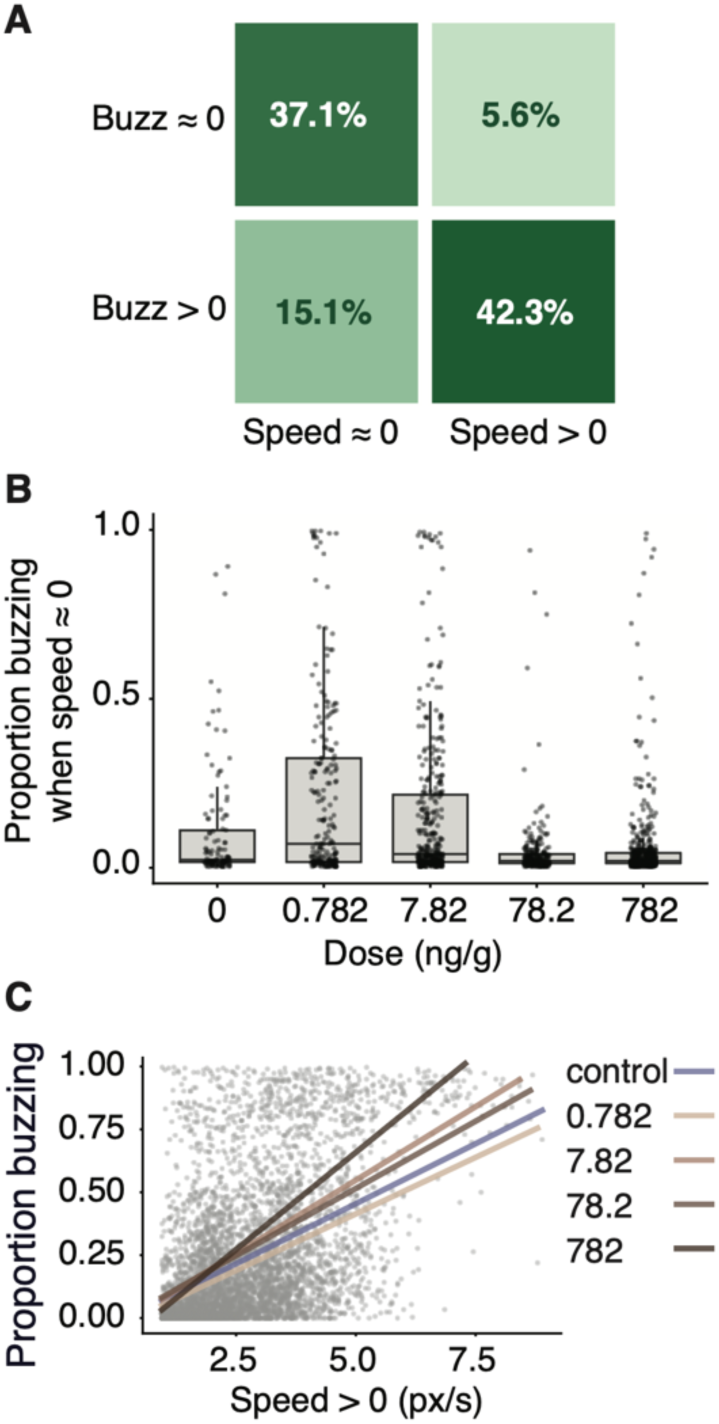
Buzzing frequently accompanies locomotion but also occurs at low locomotor activity. A: Classification of recordings (n = 10,130) according to buzzing (absent vs present) and cage-mean speed (below vs above the locomotion threshold; 0.95 px s⁻¹). Cell shading and labels show the percentage of recordings in each category. B: Proportion of time buzzing among low-speed recordings containing non-zero buzzing, by dose. Boxes show the median and interquartile range; points are individual cage-level recordings. C: Proportion of time buzzing versus cage-mean speed for recordings above the locomotion threshold. Grey points are individual cage-time recordings; lines show the fitted relationship for each dose.

When cage-mean speed was below the threshold, the proportion of time spent buzzing among recordings containing buzzing did not differ significantly with dose (Beta GLMM: χ² = 3.06, p = 0.080; Figure 4B). When bees were moving, the proportion of time buzzing increased with cage-mean speed, and this relationship varied with dose (zero-inflated Beta GLMM, speed × dose interaction: χ² = 24.9, p < 0.0001; Figure 4C). These results demonstrate that acoustic and locomotor activity overlap substantially but provide non-identical measures of behavioural state.

### Moxidectin exposure reduces bumble bee survival at higher doses

Because bees could be motionless during individual recordings without subsequently dying, transient motionlessness was distinguished from the sustained motionlessness used to estimate mortality timing (Figure S2A–B). The proportion of bees classified as motionless varied through time and across doses, demonstrating that motionlessness at a single recording was not equivalent to mortality. The survival analysis included 162 bees from 54 cages across the vehicle control and 0.782–782 ng bee⁻¹ treatments. Of the 51 deaths, 42 were assigned an estimated time from sustained motionlessness, while nine end-point deaths that could not be resolved temporally were assigned to the common final mortality time. The remaining 111 bees were right-censored at the end of the experiment.

Survival declined significantly with increasing moxidectin dose (Cox proportional-hazards mixed-effects model with cage as a frailty term; likelihood-ratio χ²₄ = 49.3, p < 0.0001, Figure 5A). Survival at the end of the experiment was 93.3% in controls and 90.9%, 77.8%, 54.5%, and 20.0% at 0.782, 7.82, 78.2, and 782 ng bee⁻¹, respectively. Relative to the control, mortality hazard was significantly elevated at 78.2 ng bee⁻¹ (hazard ratio = 8.25, 95% CI 1.83–37.1, p = 0.006) and 782 ng bee⁻¹ (hazard ratio = 33.1, 95% CI 7.52–146, p < 0.0001), but not at 0.782 or 7.82 ng bee⁻¹ (p = 0.736 and 0.105, respectively). Treating dose as a continuous predictor similarly showed a strong monotonic dose–response, with mortality hazard increasing 3.97-fold per standard deviation increase in log_10_ dose (z = 6.70, p < 0.0001). Median survival was reached only at 782 ng bee⁻¹, at approximately 43.8 h. The cage-level frailty variance was small (variance = 0.080) and not significantly different from zero (likelihood-ratio χ²₁ = 0.17, p = 0.34); a Cox model using cage-clustered robust standard errors produced equivalent results. Sucrose consumption also declined with dose (Gamma GLM, χ² = 25.7, p < 0.0001), with the first significant reduction relative to control occurring at 78.2 ng bee⁻¹ (Figure 5B).

**Figure 5:**
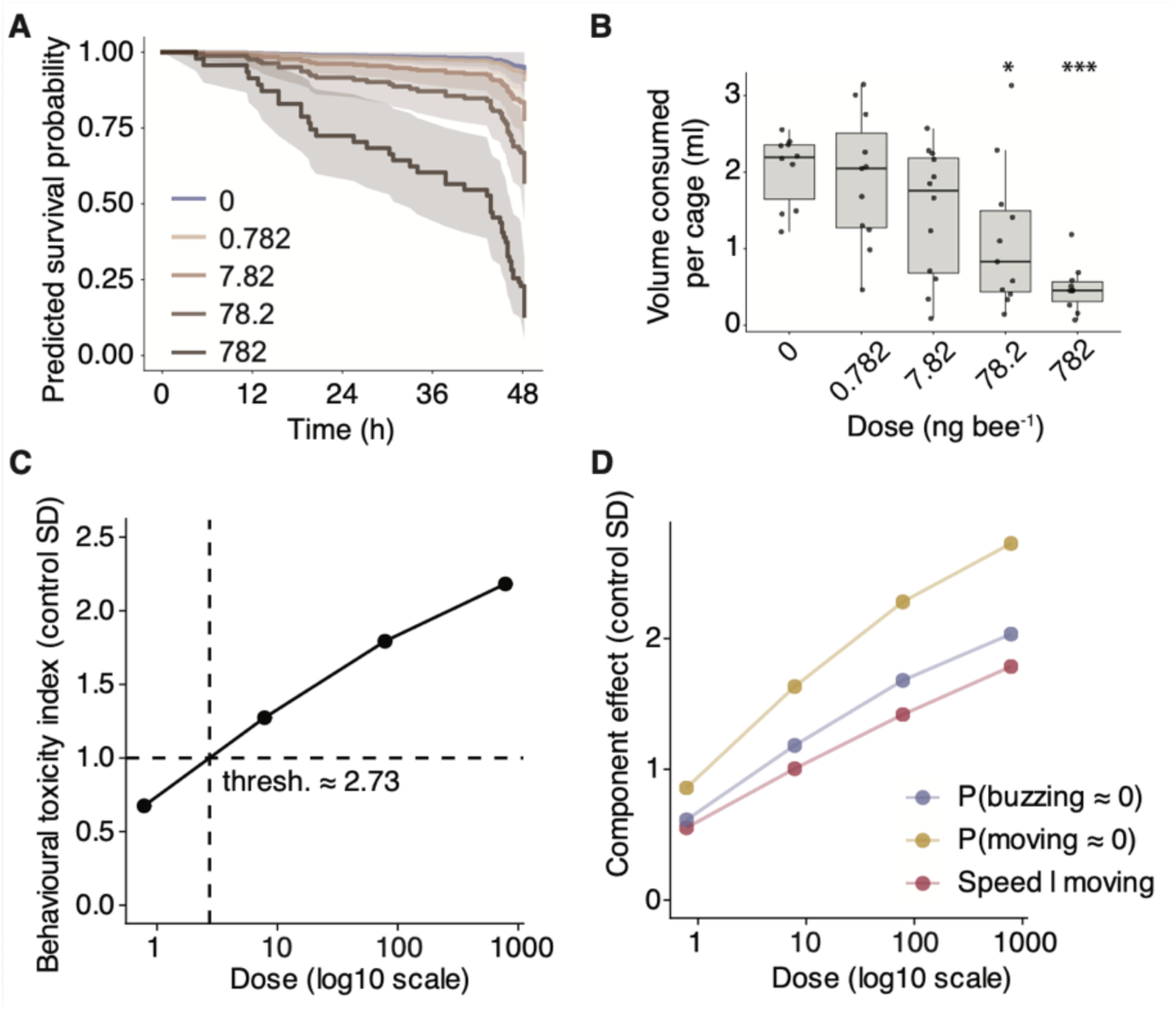
Multimodal behavioural effects occur below doses associated with increased mortality. A: Survival across doses estimated from a Cox proportional-hazards mixed-effects model with cage identity as a frailty term. Lines show model-predicted survival probabilities and shaded bands show 95% confidence intervals. B: Total sucrose solution consumption per cage (three bees per cage) across control and moxidectin treatments. Asterisks denote significant differences relative to the vehicle control based on Holm-adjusted estimated marginal means. C: Behavioural toxicity index integrating standardised deviations from control in movement probability, locomotor speed, and buzzing probability. The dashed line indicates an index value of 1, corresponding to an average behavioural deviation of one control-cage SD across the three components; the interpolated dose at which the index crossed this threshold was 2.73 ng bee⁻¹. D: Standardised dose-dependent deviations from control for each behavioural component used to construct the composite index. The components showed different dose–response patterns, with movement probability exhibiting the steepest apparent change across doses.

To determine whether mortality itself accounted for the observed reduction in locomotion, we repeated the cage-level speed analysis after excluding bees assigned as dead. Excluding dead bees increased estimated cage-level speed by approximately 9% overall, but the effect varied across doses (method × dose interaction: F_1,70_ = 7.26, p < 0.01; main effect of method: F_1,70_ = 18.41, p < 0.0001), with a significant difference between analyses only at 782 ng bee⁻¹ (Figure S3). Similarly, excluding dead bees significantly affected movement probability only at the highest dose (method × dose interaction: F_4,47_ = 3.33, p = 0.0176). Thus, mortality had little influence on estimated locomotor responses at lower doses, suggesting that the observed reductions in activity were primarily attributable to sublethal effects rather than mortality.

### Multimodal integration identifies behavioural effects below the lethal-effect range

To integrate behavioural responses across modalities, we defined a behavioural toxicity index combining dose-specific deviations from control in movement probability, locomotion speed, and buzzing probability. The resulting toxicity index increased monotonically with dose, indicating progressively larger behavioural disruption across modalities (Figure 5C). The index was expressed as the mean absolute deviation across the three behavioural components, each standardised by variability among control cages. We used an index value of 1 (an average standardised deviation of one control SD across components) as a predefined threshold for behavioural disruption. The index crossed this threshold at an estimated dose of 2.73 ng bee⁻¹ (by log-linear interpolation between tested doses), with larger deviations at higher doses. Contributions to the composite index were not uniform across components (Figure 5D), with reductions in movement probability showing the largest deviations, followed by reductions in buzzing probability, and smaller contributions from changes in locomotion speed among active bees.

## Discussion

This study demonstrates that acoustic activity provides a practical and complementary behavioural signal for detecting pesticide-induced effects in bees. Across multiple insect pollinator species, buzzing showed structured temporal variation and was often associated with locomotor activity, but the two modalities were not completely coupled: buzzing was also detected when insects showed little or no translational movement. Acoustic monitoring can therefore capture aspects of behavioural state that are not represented by locomotor measures alone. In bumble bees, both acoustic and video-derived measures showed dose-dependent responses following moxidectin exposure, while their integration identified behavioural disruption at doses substantially below those associated with significantly elevated mortality hazard. Together, the cross-species and toxicological assays demonstrate both the generalisability of BEEhaviourLab and the value of combining behavioural modalities when individual endpoints provide only a partial description of behavioural state.

Automated behavioural phenotyping has transformed toxicology and neurobiology by enabling scalable behavioural screening across model organisms including *Caenorhabditis elegans*^35^, zebrafish^10,36^, *Daphnia*^37^, and *Drosophila*^9,38,39^. Comparable systems for standardised behavioural toxicology across insect pollinators remain relatively limited. Recent advances in automated bee monitoring have begun to address this gap: in-hive and semi-field tracking systems can generate detailed behavioural datasets spanning individual and colony-level processes^40^, while laboratory platforms tracking tagged individuals within microcolonies enable detailed analysis of social behaviour^41^. These approaches have provided important insights into colony dynamics.

BEEhaviourLab complements these systems by providing a generalised platform for high-throughput behavioural phenotyping under controlled laboratory conditions. A single detector trained across four taxa performed consistently well despite differences in body size and morphology, supporting the use of a shared tracking framework. To our knowledge, BEEhaviourLab is the first automated insect phenotyping platform to integrate acoustic monitoring alongside video tracking. This adds a behavioural dimension that has largely been overlooked in automated laboratory phenotyping. In bees, buzzing is associated with diverse behavioural and physiological processes, including pollination, thermoregulation, defence, and social communication^16,17,42^, and can therefore provide information about motor and behavioural state beyond translational movement alone.

The acoustic validation nevertheless highlights an important distinction between comparative and absolute measurement. Automated estimates showed strong agreement with manually annotated recordings in their relative variation among recordings, but systematically overestimated absolute buzzing duration. False-positive detections occurred predominantly in recordings containing genuine buzzing, suggesting that the classifier tended to overestimate existing acoustic activity rather than generate entirely spurious detections. However, some false-positive detections may coincide with periods of low locomotion, potentially inflating estimates of buzzing without movement. The primary strength of the acoustic classifier is therefore its ability to resolve comparative differences among recordings, treatments and species rather than to provide exact estimates of the duration of individual acoustic behaviours. This distinction is particularly important when interpreting acoustic activity as a quantitative behavioural endpoint.

The relationship between buzzing and locomotion further demonstrates why the two modalities should be viewed as complementary rather than interchangeable. Buzzing frequently accompanied locomotion, particularly in the eusocial species, but was also detected when locomotor speed was low or insects were otherwise stationary. Such activity could arise from non-translational behaviours involving wing or thoracic muscle activity, defensive or communicative buzzing, or other stationary motor behaviours. We did not classify these behaviours individually, so their relative contributions remain unknown. Conversely, locomotion also occurred without detectable buzzing. The two modalities therefore capture overlapping but distinct components of behavioural state, with acoustic monitoring identifying activity that positional tracking alone may classify as inactivity.

Applying this multimodal framework to moxidectin exposure revealed substantial behavioural effects in Bombus terrestris. Macrocyclic lactones are widely used in livestock management, and their ecological effects have been studied primarily in dung-associated arthropods^20,28^, whereas potential effects on pollinators remain poorly understood. Increasing moxidectin dose reduced both the probability of movement and locomotor speed among active bees, and reduced acoustic activity and altered its diel organisation. Similar disruptions to locomotor, diel, and flight-related behaviour have been reported following exposure to other neuroactive pesticides, including neonicotinoids^12,13,43,44^. Because changes in basic motor activity can scale to higher-level outcomes such as altered foraging and resource collection, the impairments observed here could have broader consequences for individual and colony performance, although such outcomes were not measured directly in this study.

The acoustic response also extends previous evidence that pesticide exposure can affect bee-generated sound. Thiamethoxam alters the development of buzz-pollination sonication characteristics in *B. terrestris*^45^, while *B. impatiens* exposed to imidacloprid are less likely to produce pollination sonications^46^. These studies focused specifically on buzz-pollination behaviour, whereas our passive acoustic approach detects buzzing throughout the recording period without requiring a predefined behavioural context. This broadens the potential use of acoustic measurements as ecotoxicological endpoints, particularly where behavioural effects may not be captured by locomotor activity alone.

The observed responses also raise the possibility that veterinary pharmaceuticals represent an underappreciated source of risk to pollinating insects. Although moxidectin residues have been measured in livestock dung^20^, the topical doses used here were intended to establish dose–response relationships rather than reproduce field exposure levels. Pollinators may encounter veterinary pharmaceuticals through complex pathways that depend on environmental persistence, soil contact, nesting behaviour and other factors, and the extent of realistic exposure through these routes remains uncertain. Establishing exposure under environmentally realistic conditions will therefore be essential before the present dose–response relationships can be translated directly into field risk.

A central question was whether the observed reductions in movement could instead reflect increasing mortality. Continuous behavioural monitoring was useful here because immobility alone cannot reliably distinguish severe impairment from death. Bees could remain motionless during individual recordings and subsequently resume activity, and pesticide-exposed individuals may adopt rigid, inverted or otherwise atypical postures that further complicate visual classification^44^. By identifying sustained increases in cage-level motionlessness that were consistent with independently recorded mortality, we reconstructed approximate mortality trajectories across the experiment. Mortality hazard was significantly elevated relative to the control at 78.2 and 782 ng bee⁻¹, while median survival was reached only at 782 ng bee⁻¹. Sucrose consumption was also first significantly reduced at 78.2 ng bee⁻¹. In contrast, locomotor and acoustic responses changed progressively across lower doses, supporting the interpretation that behavioural disruption preceded the strongest conventional toxicity responses.

We additionally tested whether mortality disproportionately affected estimates of locomotor impairment by recalculating cage-level speed and movement probability after excluding bees from recordings following their inferred mortality time. This increased cage-mean locomotor speed by approximately 9% overall, but significant differences between the unadjusted and mortality-excluded analyses were observed only at the highest dose, for both speed and movement probability. The overall dose-dependent patterns were retained, suggesting that reductions in locomotor activity at lower doses were not simply a consequence of dead individuals remaining in the cages. A similar analysis was not possible for acoustic activity, as buzzing was measured at the cage level and could not be attributed to individual bees. Consequently, we cannot fully separate the effects of mortality and sublethal behavioural impairment on buzzing probability. More broadly, this illustrates an advantage of longitudinal behavioural monitoring: transient inactivity, behavioural impairment and mortality can be considered separately rather than being collapsed into a single endpoint.

Integrating the behavioural responses into a multimodal index provided a complementary way to summarise the onset of behavioural disruption. The index combined the probability of moving, locomotor speed conditional on movement, and the probability of buzzing, thereby capturing whether bees were active, how rapidly they moved when active, and whether they produced detectable acoustic activity. The three components showed different dose–response patterns, with movement probability exhibiting the steepest apparent change, but no formal comparison was made of their relative contributions. Their combination was intended to reduce dependence on any single endpoint and to represent multiple manifestations of behavioural impairment rather than to identify a single most sensitive metric.

The combined index crossed its predefined threshold, corresponding to an average deviation of one control-cage standard deviation, at an estimated dose of 2.73 ng bee⁻¹. This interpolated behavioural-effect threshold was below the lowest tested dose associated with significantly elevated mortality hazard, 78.2 ng bee⁻¹, although because our doses were spaced an order of magnitude apart, the separation between behavioural and lethal thresholds cannot be quantified precisely. The index should be interpreted as a summary measure rather than as a universally validated toxicity threshold. We have not tested whether the same combination of endpoints is optimal for other chemical stressors, nor whether an acoustic-only or locomotor-only index would produce an equivalent dose–response relationship. Testing the transferability and robustness of multimodal indices across compounds with different modes of action will therefore be an important next step.

Several additional features of the study constrain interpretation. Tag-free tracking does not provide independently verified biological identities, and trajectories were linked using frame-to-frame positional continuity. Identity exchanges may therefore occur when insects cross or overlap. Because treatment-level locomotor analyses were aggregated within cage, persistent exchanges of numerical trajectory labels do not alter the cage-level quantities used for inference. Short-lived correspondence errors could nevertheless introduce noise into frame-to-frame displacement estimates and potentially inflate some instantaneous speed measurements. Similarly, carrying positions forward across detection gaps rather than interpolating movement may introduce bias into derived motion metrics.

Acoustic measurements have related interpretative limitations. Although the classifier detected most manually annotated buzzing events, its lower precision and systematic overestimation of buzzing duration mean that changes in acoustic activity should be interpreted primarily as relative differences in detectable buzzing-related sound. The classifier did not identify specific behavioural categories, and the present data therefore cannot establish whether changes arose from communication, thermoregulation, defence, flight-related activity or other sound-producing behaviours. Future analyses that combine acoustic features with posture and movement classification may be able to resolve these behavioural states more directly.

Cross-species comparisons should likewise be interpreted cautiously. Bumble bees were laboratory-reared whereas the other species were wild-caught, creating differences in rearing and entrainment history that could influence diel patterns. All insects in the comparative assay were also housed singly to standardise recording conditions. While this approximates the natural condition of solitary species, social isolation may itself alter behaviour in *A. mellifera* and *B. terrestris*. Their greater acoustic activity could therefore partly reflect isolation rather than intrinsic species differences, although genuine differences in the behavioural use of acoustic and vibrational signals may also contribute^16,17,42^. Species-level inference is most limited for *C. hederae* and *E. tenax*, which were sampled less extensively than the other taxa. Their contrasting buzzing profiles compounded this: *C. hederae* was frequently silent (21.3% of recordings contained no detected buzzing), whereas *E. tenax* buzzed briefly in every recording, leaving its zero-inflation estimate unidentifiable.

Overall, this study shows that combining acoustic monitoring with automated video tracking can reveal behavioural effects that would be only partially represented by either modality alone. BEEhaviourLab provides a scalable framework for quantifying behavioural impairment across insect taxa while linking sublethal behavioural responses to conventional toxicity endpoints. Continuous trajectory and audio data also create opportunities for more detailed behaviour segmentation and state-based modelling, including automated classification of feeding, resting, grooming and abnormal motor activity. Incorporating such complementary behavioural measures into ecotoxicological screening could help identify chemical effects before they are evident through mortality alone and provide a stronger mechanistic bridge between laboratory toxicity assays and the ecological consequences of environmental exposure.

## Methods

### BEEhaviourLab equipment

BEEhaviourLab is a modular, distributed platform for automated, high-throughput recording of insect behaviour under controlled experimental conditions. The system can scale from a single recording unit to many parallel nodes, each operating independently but coordinated through a central control computer (Figure 1A–B). Each node integrates synchronised video recording (Raspberry Pi NoIR v2 camera), audio capture (RØDE smartLav+ microphone), and environmental monitoring (temperature and humidity via a DHT22 sensor mounted within each recording arena). All experiments were performed at room temperature (23.7 °C; 95% CI: 22.1 – 24.9 °C). Illumination was provided beneath the transparent Perspex recording arena using LED strip lights, which operated in white (daylight) or red (night-mode) settings. Red illumination was used at night because bees and flies generally have reduced visual sensitivity at longer wavelengths, thereby minimising visual stimulation while permitting video recording^47,48^. The physical enclosure was constructed from laser-cut Perspex and assembled using plastic fasteners or adhesive (Figure 1A). A Raspberry Pi 4B single-board computer (4 GB RAM) managed recording and sensor acquisition.

Experimental parameters were defined in a human-readable configuration file, allowing users to specify metadata (e.g. species, treatment, replicate), recording duration, inter-recording intervals, number of recording segments, and video and audio capture settings (Figure 1C). This approach enabled identical experimental protocols to be deployed across multiple nodes with minimal manual intervention. Nodes were accessed remotely via SSH over Wi-Fi from the central control computer, which initiated recordings and coordinated data acquisition across the system. Each recording segment was automatically compressed and stored with structured metadata before being transferred to the central computer for storage and downstream analysis.

All hardware designs and Raspberry Pi control software are fully open source and available on GitHub (BEEhaviourLab/BEEhaviourLab-apparatus). The modular architecture allows users to adapt the system for different experimental needs, including adding sensors, modifying acquisition parameters, or extending recording and control functionality.

### Automated insect tracking

Insects in BEEhaviourLab video recordings were automatically detected and tracked using YOLO version 8 “nano” (YOLOv8n, Ultralytics), a lightweight object-detection architecture suitable for rapid fine-tuning and deployment on standard computing hardware. To support reuse and adaptation, we provide a step-by-step tutorial for retraining the model on user-defined insect datasets (GitHub: BEEhaviourLab/BEEhaviourLab-YOLO-training). This repository also contains the fine-tuned models reported in this study.

Training and validation data were derived from video frames extracted from BEEhaviourLab recordings. Frames were manually annotated in Label Studio (labelstud.io) with two object classes: *bee* and *feeder*. Two YOLOv8n models were trained for 100 epochs: model 1 was trained on 155 annotated images from recordings of bumble bees (*Bombus terrestris*); and model 2 was trained on a multi-species dataset comprising 455 annotated images across four insect species: honey bee (*Apis mellifera*), ivy bee (*Colletes hederae*), bumble bee (*Bombus terrestris*), and hoverfly (*Eristalis tenax*).

Model performance was evaluated on validation datasets consisting of recordings from independent experimental cohorts, ensuring no overlap between training and evaluation data. Performance metrics included recall (proportion of true objects detected), precision (proportion of detections that are correct), mAP50 (mean average precision calculated using detections with at least 50% overlap between predicted and true bounding boxes), mAP50-95 (mean average precision averaged across overlap thresholds from 50% to 95%), and the F1-score (harmonic mean of precision and recall). Following model development, an independent test dataset was used to evaluate detector performance. The trained model and hyperparameters were fixed before evaluation on this test set. Test images (20 honey bee and 20 bumble bee) were sampled from recordings that were not represented in either the training or validation datasets and were manually annotated using the same procedure as the training data.

Initial tracking across frames was performed using the BoT-SORT algorithm^49^, which is designed for scenarios where the number of tracked objects may vary over time. The algorithm includes mechanisms to recover identities when detections are temporarily lost (e.g. due to occlusion), but in our recordings it frequently generated new IDs when individuals reappeared. Because the number of insects in the arena was fixed, we applied an additional post-processing step after BoT-SORT tracking. For each frame, detections were matched to a fixed set of identities (IDs 1…N) using the Hungarian algorithm^50^, minimising Euclidean displacement from positions in the previous frame. When detections were temporarily missing, the last known position was propagated forward while the corresponding detection was recorded as missing, maintaining constant population size and preventing the creation or deletion of identities. These IDs represent trajectory labels rather than independently verified biological identities, and labels may occasionally be exchanged when individuals cross or overlap.

Locomotor activity was quantified from the tracked positions of each individual insect across frames. Pixel coordinates were converted to physical distance using a calibration factor derived from the known dimensions of the recording arena, allowing changes in position to be expressed in mm regardless of recording resolution. Movement speed (mm s^-1^) was calculated as the frame-to-frame displacement of each insect multiplied by the video frame rate (30 frames s^-1^) and used as a simple, robust measure of activity for comparing behavioural responses across treatments and time periods.

### Audio analyses

Audio recordings were processed using a custom pipeline to detect segments containing buzzing-related acoustic activity. Recordings were resampled to 16 kHz and bandpass-filtered (200–7920 Hz) to focus on bee-relevant frequencies. The filtered signal was divided into 1 s windows with a 0.2 s hop, and frame energy was calculated in decibels as

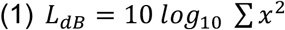

A robust threshold based on the median frame energy and its normalised median absolute deviation (MAD) was used to classify frames as containing sound or silence. MAD was calculated as:

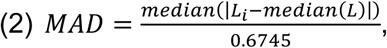

where the scaling factor expresses MAD on a standard-deviation-equivalent scale for normally distributed data. For recordings dominated by background noise, frames exceeding

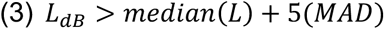

were classified as containing sound. For recordings whose median frame energy was > −30 dB (53.6% of files), the recording was treated as signal-dominated and a lower threshold was used (the minimum of the 10th percentile, 25th percentile, and [median − 3 × MAD] of frame energy, further lowered to the 5th percentile or [median − 4 × MAD] when median energy exceeded −25 dB, and floored just above the recording minimum) so that predominantly active files were not thresholded as if the median were background. Frames were labelled as sound or silence every 0.2 s. Rules to drop very short sounds (<0.1 s) or to bridge very short silences (<0.15 s) were included in the code, but both are shorter than the 0.2 s step between frames, so they never changed the labels. For each 60 s audio segment, the proportion of time containing detected buzzing-related acoustic activity was the proportion of these 1 s frames classified as sound.

Automated buzzing classification was validated against manual annotation of fifty 60 s recordings (maximum one per cage replicate). Thirty-five recordings were of *Bombus terrestris* from the moxidectin experiment, stratified across dose (0–782 ng bee⁻¹) and day/night. The remaining 15 were from the multi-species dataset (5 each of *<u>Apis mellifera</u>*, *Colletes hederae* and *Eristalis tenax*). Recordings were further stratified using a coarse automated activity class (no buzzing, some buzzing, or extensive buzzing) so that both silent and active recordings were represented; these automated scores were not shown to the annotator. Each clip was anonymised and scored. An observer inspected the waveform and spectrogram in Audacity while listening and marked onset–offset intervals containing buzzing from one or more bees. Unlabelled time was treated as no buzzing sound.

Manual intervals were converted onto the same analysis frames as the detector (1.0 s energy windows, 0.2 s hop; 296 frames per 60 s clip). A frame was classed as buzzing if labelled sound occupied at least 50% of the window. The detector was re-run on the annotated clips with the same parameters used in the main analysis. Frame-level agreement was summarised with precision, recall, F1-score, specificity and balanced accuracy (silence was the majority class). At clip level, automated proportion of time buzzing was compared with the labelled duration divided by 60 s, using Pearson and Spearman correlation and mean absolute error.

### Cross-species behavioural recordings

To evaluate the flexibility of the automated tracking pipeline and quantify locomotor and acoustic activity across taxa, four insect species were recorded in BEEhaviourLab: honey bee (*Apis mellifera*, n = 21), ivy bee (*Colletes hederae*, n = 3), hoverfly (*Eristalis tenax*, n = 5), and bumble bee (*Bombus terrestris*, n = 33). Honey bees, ivy bees, and hoverflies were wild-caught immediately prior to recording and released afterwards. Bumble bees were obtained from commercial colonies (Biobest, Westerlo, Belgium; via Agralan, Swindon, UK) and kept at room temperature with ad libitum access to sugar syrup and honey bee-collected pollen three times per week. Wild insects were caught by net while foraging on flowers, transported in plastic vials, and transferred to a BEEhaviourLab cage within 2 h of capture. We included only female insects (worker caste for honey bees and bumble bees) in the assays, identifying them visually by body size and eye morphology. Insects were housed one per cage in the BEEhaviourLab.

Recordings were conducted under a 12 h light:12 h dark cycle (lights on 06:00–18:00), and data were collected from late August to mid-September 2024 (actual day length approximately 14 h). Video and audio recordings consisted of 60 s segments separated by 29 min intervals over a 48 h period, yielding 96 recordings per cage. These data were used to evaluate detection performance and quantify locomotor and acoustic activity across species.

### Moxidectin toxicity in bumble bees

To demonstrate the application of BEEhaviourLab for sublethal toxicity assessment, we quantified behavioural effects of moxidectin on *Bombus terrestris* workers over a 48 h period following contact exposure. Commercial bumble bee colonies were kept as described above. Individual bees received a 1 µl droplet applied to the dorsal thorax containing moxidectin (Cydectin Pour-On formulation, 0.5% w/v) diluted in DMSO. Exposure doses were 0 (vehicle control; n = 30), 0.782 (n = 33), 7.82 (n = 36), 78.2 (n = 33), and 782 (n = 30) ng bee⁻¹. We included bees from three colonies, balanced across treatments. The dose series spanned several orders of magnitude, with the upper dose numerically corresponding to the maximum reported moxidectin concentration in cattle dung of 782 ng g⁻¹, which remained at 490 ng g⁻¹ after 42 days of field exposure20. However, because dung concentrations cannot be directly converted into doses received by individual bees, these values were used as a reference for selecting a broad experimental dose range rather than to estimate environmentally realistic exposure. Field measurements of moxidectin in soil and estimates of uptake by wild bees remain unavailable.

Following exposure, bees were placed in clear Perspex recording cages with 3 bees per cage (10 control cages, 11 cages at 0.782 ng bee⁻¹, 12 at 7.82 ng bee⁻¹, 11 at 78.2 ng bee⁻¹, and 10 at 782 ng bee⁻¹) and provided with 1 M sucrose via a syringe feeder. Feeders were weighed before and after the experiment, and the mass consumed was converted to volume using the density of the sucrose solution, with losses due to evaporation corrected using feeders placed in empty cages over the same period.

Recordings were conducted under a 12 h light:12 h dark cycle (lights on 06:00–18:00) and data were collected in June 2025 (actual day length ∼16.5 h). Video and audio were recorded for 60 s, separated by 14 min intervals over a 48 h period, yielding 192 recordings per cage. At the end of the experiment, sucrose consumption, body mass, and cage-level mortality were recorded. Surviving bees were euthanised by freezing overnight.

To estimate the timing of mortality from the video recordings, tracked bees were classified as motionless at each recording time point when their maximum frame-to-frame displacement remained below a predefined threshold (<7 pixels per frame). The threshold was manually validated by inspecting known motionless bees and identifying the maximum frame-to-frame displacement attributable to tracking noise, including variation in YOLO detection centroid position. Because individual identities could not be maintained reliably across successive recordings, mortality timing was inferred at the cage level from changes in the number of persistently motionless bees. For a cage in which *k* deaths were recorded by the end of the experiment, the *r*th death was assigned to the earliest recording at which at least *r* bees were motionless and remained so in all subsequent recordings. Motionless counts were capped at the total number of deaths recorded for that cage.

### Statistical analyses

Statistical analyses were performed in R version 4.5.2^51^. A summary of models is provided in Table S1. Behavioural data were analysed using generalised linear mixed-effects models implemented in `glmmTMB`^52^. For repeated behavioural measurements, cage identity was included as a random intercept to account for repeated observations of the same group through time. In the moxidectin experiment, three bees were housed and recorded together in each cage and individual identities were not maintained across recording periods; behavioural responses were therefore analysed at the cage level rather than treating individual bees as independent repeated measures. Model diagnostics were assessed using simulation-based residual checks in `DHARMa`^53^ and model-based estimates and contrasts were obtained using `emmeans`^54^. Model predictions and effect sizes are reported with 95% confidence intervals derived from the fitted mixed-effects models using Wald approximations based on the estimated standard errors.

Locomotor behaviour was analysed using a two-part mixed-effects framework separating behavioural state from movement intensity. The bumble bee speed distribution was strongly bimodal on a logarithmic scale, with distinct inactive and moving components. The movement threshold was placed in the trough separating these components, which is bracketed by two independent estimates: the antimode of the kernel density of log₁₀ speed (0.58 px s⁻¹) and the equal-posterior boundary of a two-component Gaussian mixture fitted to log₁₀ speed (1.63 pixels s⁻¹). We adopted 0.95 px s⁻¹, the approximate geometric midpoint of these two estimates (Figure S1). The threshold was calculated once from the pooled bumble bee dataset and applied consistently across all treatments and recording times. Activity classification was performed using pixel-based speed, while speeds used for quantitative locomotor comparisons were converted to mm s^-1^ using the arena calibration described above. Movement probability was modelled using a binomial GLMM with a logit link, while movement speed conditional on activity was modelled using a Gamma GLMM with a log link.

For the moxidectin experiment, locomotor data were aggregated within cage at each time point before modelling. Movement probability was represented as binomial counts of moving versus inactive bees within each three-bee cage, while movement speed conditional on activity was calculated as the mean speed of active bees within each cage. For these models, moxidectin concentrations (transformed with log_10_[dose + 0.0782]) and time (z-scored) were included as covariates. For the cross-species dataset, species and time were included as fixed effects. Audio-derived behavioural measures were analysed using zero-inflated beta GLMMs, separating the probability of zero buzzing activity from the distribution of non-zero proportions, with the same dose and time covariates as for the locomotion data. Diel patterns of activity were analysed using harmonic mixed-effects models including 24 h sine and cosine terms. The presence of diel rhythmicity and its modulation by treatment or species were evaluated using likelihood-ratio tests.

We tested the impact of removing bees that were assigned as dead due to continued motionlessness on cage-level locomotor speed and movement probability, because the primary analysis retained these individuals when calculating per-cage values. Movement probability was re-analysed using the binomial GLMM described above. Cage-mean walking speed was modelled with a Gamma generalised linear mixed model (log link), matching the primary locomotion-speed analysis. Fixed effects were scaled dose (log_10_[dose + 0.0782]), scaled time (z-scored) since the start of recording, and a two-level factor indicating whether bees assigned as dead were included or excluded from the cage mean. A random intercept was included for cage. For the dead-excluded dataset, cage-mean speed was the sum of track speeds divided by the number of remaining live tracked bees (*n*_existing_ − *n*_dead_). Estimated marginal means were obtained at the experimental doses. Treatments were compared with control within each dataset, and the two datasets were compared at each dose, with Holm adjustment.

Survival following moxidectin exposure was analysed using Cox proportional-hazards mixed-effects models (`coxme`), with each bee treated as an individual subject and cage included as a frailty term to account for the three bees housed together. Dose was fitted in separate models as a five-level factor, yielding hazard ratios relative to the vehicle control, and as a continuous predictor to test for a monotonic dose–response trend. Deaths identified from sustained motionlessness were assigned their estimated times as described above. Where end-point mortality could not be matched to a sustained transition in the video record, death was conservatively assigned to 2,895 min, 15 min after the final recording; surviving bees were censored at the same time. Two cages with prematurely terminated recordings had no recorded deaths and were likewise followed to 2,895 min. The overall effect of dose was assessed by likelihood-ratio test against a model containing only the cage frailty term, and individual doses were compared with control using Wald tests. The contribution of cage was assessed by a boundary-corrected likelihood-ratio test, and results were confirmed using a Cox model with cage-clustered robust standard errors. The proportional-hazards assumption was assessed using scaled Schoenfeld residuals (`cox.zph`), and model-predicted survival curves were generated for each dose.

To examine the relationship between acoustic and locomotor activity, each recording was classified by whether buzzing was absent or present (proportion buzzing = 0 vs > 0) and whether cage-mean speed was below or above the locomotion threshold defined above (0.95 px s^-1^). For recordings below this threshold with non-zero buzzing, the proportion of time buzzing was modelled with a Beta GLMM (`glmmTMB`, logit link; n = 1,525 recordings, 54 cages). For recordings above the threshold, the proportion of time buzzing was modelled with a zero-inflated Beta GLMM (`glmmTMB`, logit link; n = 4,846 recordings, 54 cages). The conditional (Beta) component included cage-mean speed, dose, and their interaction as fixed effects, with cage identity as a random intercept; the zero-inflation component included speed and dose as fixed effects.

To summarise multimodal behavioural effects, we calculated a behavioural toxicity index from cycle-averaged, cage-level predicted deviations from control behaviour in movement probability, locomotion speed, and buzzing probability. For each metric, the absolute deviation from the control prediction was divided by the standard deviation (SD) among control cages, expressing effects in units of natural control variability. The three standardised effects were then averaged to obtain a single index for each dose. An index value of 1 therefore represents an average behavioural deviation of one control-cage SD, and this threshold was used to estimate the dose at which sublethal behavioural effects first emerged. Cage-level sucrose volume was modelled with a Gamma GLM (log link). Treatments were compared with control using Holm-adjusted estimated marginal means (`emmeans`).

## Author contributions

Conceptualization: R.H.P. Methodology: R.H.P., J.S., O.N.F.K., and C.N. Software: R.H.P., O.N.F.K., and C.N. Investigation: J.C.Z.K., A.S., and K.W. Data curation: R.H.P. and O.N.F.K. Formal analysis: R.H.P. and O.N.F.K. Visualization: R.H.P. Supervision: R.H.P., G.A.W., and S.R. Project administration: R.H.P. Funding acquisition: R.H.P. Writing (original draft): R.H.P. Writing (review and editing): All authors.

## Acknowledgements

Not applicable.

## Data Availability

All data used in the analyses shown in this paper are available online (https://doi.org/10.6084/m9.figshare.31465879).

## Code Availability

All code is available through the BEEhaviourLab GitHub page. Files for constructing the BEEhaviourLab apparatus: github.com/beehaviourlab/beehaviourlab-apparatus. Python code and tutorials for training a new YOLO model and testing the models in this manuscript: github.com/beehaviourlab/beehaviourlab-yolo-training. The main BEEhaviourLab repository with code for analysing the video and audio recordings and generating time-series plots: github.com/beehaviourlab/beehaviourlab. The R code for the statistics used in this paper can be found at https://doi.org/10.6084/m9.figshare.31465879.

