## Supplemental information for "Acoustic phenotyping reveals sublethal pesticide effects in bees using BEEhaviourLab"

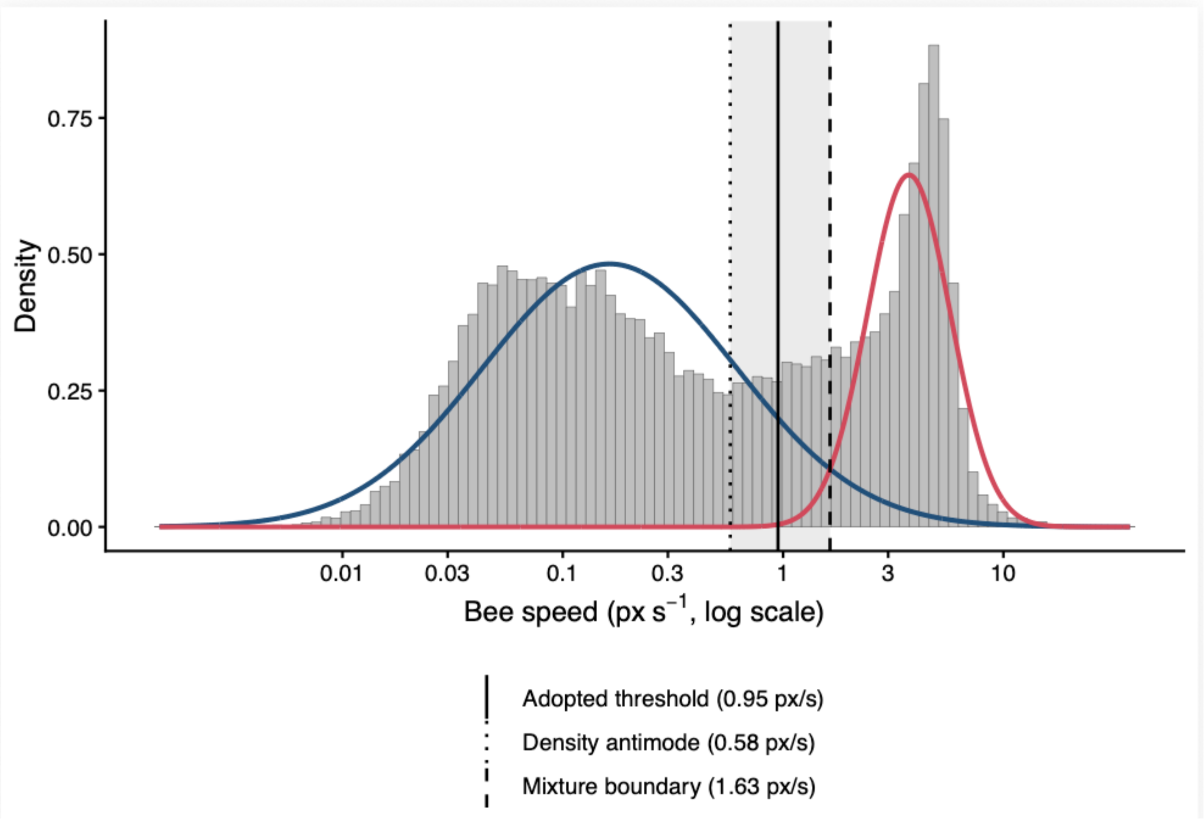


**Figure S1: Derivation of the locomotion speed threshold.**

Distribution of individual bee speeds across all cages, timepoints and doses (*n* = 28,366 non-zero observations), shown on a logarithmic scale. The distribution is bimodal, corresponding to inactive and actively moving bees. Curves show a two-component Gaussian mixture fitted to $\log_{10}$ speed: an inactive component containing 69.5% of observations and centred at 0.16 px s$^{-1}$ (blue), and a moving component containing 30.5% and centred at 3.73 px s$^{-1}$ (red). Vertical lines indicate the adopted movement threshold (0.95 px s$^{-1}$, solid) based on two independent data-driven estimates of the boundary between components: the kernel-density antimode (0.58 px s$^{-1}$, dotted) and the mixture equal-posterior boundary (1.63 px s$^{-1}$, dashed). The adopted threshold falls at the approximate midpoint of these two estimates.

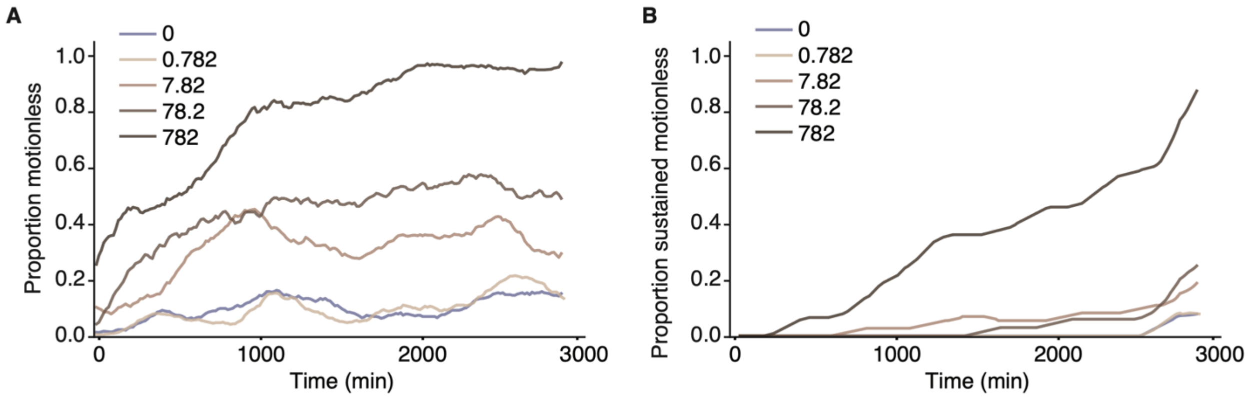


**Figure S2: Instantaneous and sustained motionless bees over time by moxidectin dose.**

A: Bees were classified as motionless within a recording if their maximum frame-to-frame displacement was <7 pixels per frame. This classification includes transiently motionless bees that subsequently resumed activity. Curves show the proportion of bees classified as motionless at each recording, pooled by dose and smoothed with a 15-point centred rolling mean. Colours indicate moxidectin dose (0, 0.782, 7.82, 78.2, and 782 ng $bee^{-1}$). B: Proportion of bees classified as persistently motionless from each recording through the end of the experiment, calculated as the future running minimum of the cage-level motionless count. This sustained-motionlessness criterion was used to estimate mortality timing for bees confirmed dead at the end of the experiment.

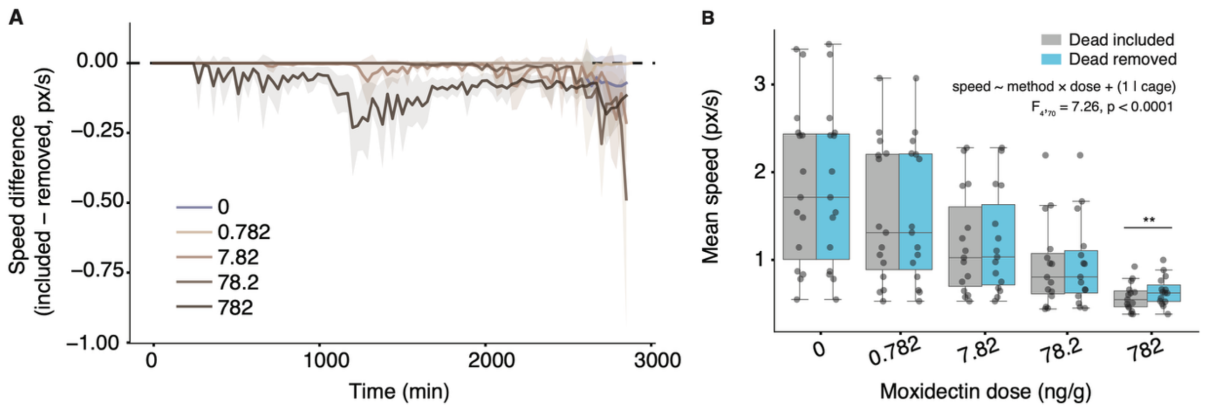


**Figure S3: Effect of excluding bees assigned as dead on cage-level locomotor speed.**

A: Change in cage-mean locomotor speed after excluding bees assigned as dead based on sustained motionlessness. Cage-level speed was calculated at each time point in two ways: the unadjusted mean (total tracked speed divided by the number of bees initially present in the cage) and the dead-excluded mean (total speed of bees not yet assigned as dead divided by the number remaining alive according to the estimated mortality times). The plotted quantity is the difference (unadjusted speed minus dead-excluded speed); negative values therefore mean that retaining bees assigned as dead reduced the estimated cage-mean speed. B: Distribution of unadjusted and dead-excluded cage-mean locomotor speeds within each dose. Boxes show the median and interquartile range (IQR) and whiskers extend to 1.5 × IQR. Asterisks indicate significant differences between unadjusted and dead-excluded speeds within a dose based on the mixed-effects sensitivity analysis.

**Table S1: Response variables, predictors, and model structure for each statistical test.** Cage (ID) was included as a random effect across models. Time and locomotion speed were z-scored, and moxidectin concentrations were scaled using log$_{10}$[dose + 0.0782].

<table>
<tr><th>Fig</th><th>Analysis</th><th>Model</th><th>Formula</th></tr>
<tr><td>2B</td><td>P(speed > threshold)</td><td>Binomial GLMM, logit</td><td>is_moving ~ Species + zscored(time) + (1 | ID)</td></tr>
<tr><td>2C</td><td>speed | speed > threshold</td><td>Gamma GLMM, log link</td><td>speed ~ Species + zscored(time) + (1 | ID)</td></tr>
<tr><td>2H</td><td>P(buzzing = 0)</td><td>Zero-inflated Beta GLMM, logit (ZI part)</td><td>ziformula = ~ Species on prop_time</td></tr>
<tr><td>2I</td><td>proportion buzzing | buzzing > 0</td><td>Same ZI-Beta GLMM, logit (conditional Beta part)</td><td>prop_time ~ Species + zscored(time) + (1 | ID)</td></tr>
<tr><td>3B</td><td>P(speed > threshold)*</td><td>Binomial GLMM, logit</td><td>is_moving ~ log$_{10}$[dose + 0.0782] + zscored(time) + (1 | ID)</td></tr>
<tr><td>3C</td><td>speed | speed > threshold</td><td>Gamma GLMM, log link</td><td>speed ~ log$_{10}$[dose + 0.0782] + zscored(time) + (1 | ID)</td></tr>
<tr><td>3E</td><td>P(buzzing = 0)</td><td>Zero-inflated Beta GLMM, logit (ZI part)</td><td>ziformula = ~ log$_{10}$[dose + 0.0782]on prop_time</td></tr>
<tr><td>3F</td><td>proportion buzzing | buzzing > 0</td><td>Same ZI-Beta GLMM, logit (conditional Beta part)</td><td>prop_time ~ log$_{10}$[dose + 0.0782] + zscored(time) + (1 | ID)</td></tr>
<tr><td>4B</td><td>Proportion buzzing | speed < threshold</td><td>Beta GLMM, logit</td><td>prop_buzzing ~ zscored(speed) * log$_{10}$[dose + 0.0782] + (1 | ID)</td></tr>
<tr><td>4C</td><td>Proportion buzzing | speed > threshold</td><td>Zero-inflated Beta GLMM, logit (ZI part)</td><td>prop_buzzing ~ zscored(speed) + log$_{10}$[dose + 0.0782] + (1 | ID)</td></tr>
<tr><td>5A</td><td>Survival (mortality)</td><td>Cox proportional hazards model</td><td>Surv(time, event) ~ Dose_factor + (1 | ID)</td></tr>
<tr><td>5B</td><td>Volume consumed</td><td>Gamma GLM, log link (no random effect)</td><td>vol_total ~ log$_{10}$[dose + 0.0782] + Colony + Date</td></tr>
</table>
